# *Enterococcus faecalis* modulates immune activation and slows healing during wound infection

**DOI:** 10.1101/125252

**Authors:** Kelvin Kian Long Chong, Wei Hong Tay, Baptiste Janela, Mei Hui Adeline Yong, Tze Horng Liew, Leigh Madden, Damien Keogh, Timothy Mark Sebastian Barkham, Florent Ginhoux, David Laurence Becker, Kimberly A. Kline

**Author notes:** These authors contributed equally to this work (Kelvin Kian Long Chong, Wei Hong Tay).

## Abstract

*Enterococcus faecalis* is one of most frequently isolated bacterial species in wounds yet little is known about its pathogenic mechanisms in this setting. Here, we used a mouse wound excisional model to characterize the infection dynamics of *E. faecalis* and show that infected wounds result in two different states depending on the initial inoculum. Low dose inocula were associated with short term, low titer colonization whereas high dose inocula were associated with acute bacterial replication and long term persistence. High dose infection and persistence were also associated with immune cell infiltration, despite suppression of some inflammatory cytokines and delayed wound healing. During high dose infection, the multiple peptide resistance factor (MprF) which is involved in resisting immune clearance, contributes to *E. faecalis* fitness. These results comprehensively describe a mouse model for investigating *E. faecalis* wound infection determinants, and suggest that both immune modulation and resistance contribute to persistent, non-healing wounds.

## INTRODUCTION

Wound infections affect between 7-15% of hospitalized people globally [1]. *E. faecalis* is one of the most frequently isolated bacterial species across all types of wounds, including diabetic foot ulcers, burns, and surgical sites [2–4]. In surgical site infections (SSIs), *E. faecalis* is the third most commonly isolated organism [5, 6]. *E. faecalis* infections are increasingly difficult to treat due to their intrinsic and acquired resistance to a range of antibiotics [7]. Despite the high frequency of *E. faecalis* in wound infections, little is known about its pathogenic strategies in this niche.

Bacterial biofilms, which are often polymicrobial in nature, are a major factor in wound healing [8–10]. Biofilm-associated wound infections are associated with a poorer prognosis [8, 9]. Moreover, biofilm formation promotes survival and persistence of infecting microbial species because it facilitates defence against the host immune response [11]. *E. faecalis* encodes several factors that contribute to biofilm formation, including two sortase enzymes, SrtC and SrtA, that polymerize and attach endocarditis- and biofilm-associated pili (Ebp) to the cell wall, respectively [12–14]. Ebp pili aid in the attachment of *E. faecalis* to surfaces, which is required in the initial stages of biofilm formation *in vitro* and *in vivo* during catheter-associated urinary tract infection (CAUTI) [15, 16]. Other biofilm-associated factors that are attached to the cell wall by SrtA include Ace, aggregation substance, and Esp [17–20].

In addition to initial adhesion and colonization, *E. faecalis* must also overcome host defences to establish infection. *E. faecalis* can modulate and evade the host immune response in a number of settings [21–24]. Biofilm formation, along with expression of the SrtA substrate aggregation substance, can promote *E. faecalis* survival within macrophages and polymorphonuclear leukocytes [25, 26]. The multiple peptide resistance factor (MprF) protein of *E. faecalis* confers resistance to antimicrobial peptides via electrostatic repulsion [27, 28], and is important for surviving both neutrophil-mediated clearance and within epithelial cells and macrophages by a variety of Gram-positive bacteria [29–31].

Previously, a mouse wound excisional model was developed to study wound healing processes [32–34]. This model has been used to examine bacterial factors required for wound infection by *Pseudomonas aeruginosa, Acinetobacter baumannii*, and *Staphylococcus aureus* [35–38]. In the current study, we characterized the dynamics of *E. faecalis* infection in the murine wound excisional model. We establish the minimal doses of *E. faecalis* required for colonization and infection of wounds. We also demonstrate a role for the innate immune defense factor MprF in wound infection, and show that modulation of early inflammatory responses and delayed wound healing are coupled with persistence of *E. faecalis* within wounds.

## MATERIALS AND METHODS

### Bacterial strains and growth conditions

Strains used are shown in Supplementary Table 1. For mouse infections, *E. faecalis* were grown statically at 37°C for 15 to 18 hours in Brain Heart Infusion (BHI) medium (Neogen, Lansing, USA) in the absence of antibiotics. Clinical strains isolated from patient wounds were provided by Tan Tock Seng Hospital, Singapore.

### Genetic manipulation

Construction of *E. faecalis* OG1RF Δ*mprF1* and Δ*mprF2* were previously described [27]. OG1RF Δ*mprF1/2* was made by subcloning the Δ*mprF1* deletion construct from pJRS213-Δ*mprF1* into pGCP213 to create pGCP213-Δ*mprF1* and then transforming the plasmid into OG1RF Δ*mprF2*. Chromosomal deletions were selected for and screened as described previously. Mutants Δ*ebpABC* and Δ*srtAC* are listed in Supplementary Table 1.

### Mouse wound excisional model

Mouse wound infections were modified from a previous study [39]. Male wild-type C57BL/6 mice (7-8 weeks old, 22 to 25g; InVivos, Singapore) were anesthetized with 3% isoflurane and the dorsal hair trimmed. Following trimming, Nair™ cream (Church and Dwight Co, Charles Ewing Boulevard, USA) was applied and the fine hair removed via shaving with a scalpel. The skin was then disinfected with 70% ethanol. A 6-mm biopsy punch (Integra Miltex, New York, USA) was used to create a full-thickness wound and 10 μl of the respective bacteria inoculum applied. The wound site was then sealed with a transparent dressing (Tegaderm™ 3M, St Paul Minnesota, USA). At the indicated time points, mice were euthanized and a 1 cm by 1 cm squared piece of skin surrounding the wound site was excised and collected in sterile 1X PBS. Skin samples were homogenized and the viable bacteria enumerated by plating onto both BHI plates and antibiotic selection plates to ensure all recovered colony forming units (CFU) correspond to the inoculating strain.

### Histology

Wound tissues were excised as described above and fixed in 4% paraformaldehyde in 1× PBS (pH 7.2) for 24 hours at 4°C. Samples were then submerged in 15% and 30% sucrose gradient for 24 hours each, embedded in Optimal Cutting Temperature (OCT) embedding media (Sakura, California, USA), and frozen in liquid nitrogen. 10 μm thin sections were then obtained with a Leica CM1860 UV cryostat (Leica Biosystems, Ernst-Leitz Strasse, Germany) and stained with hemotoxylin and eosin (H&E). Images of H&E sections were acquired using an Axio Scan.Z1 slide scanner (Carl Zeiss, Göttingen, Germany) fitted with a 20× Apochrome objective.

### Gene probe and Fluorescence *in-situ* Hybridization (FISH)

Detection of *E. faecalis* was achieved with the oligonucleotide probe 5’-GGT GTT GTT AGC ATT TCG/Cy3/-3’ (IDT Technologies, Iowa, United States). The general oligonucleotide probe 5’- GCT GCC TCC CGT AGG AGT/Alexa Fluor^®^ 488/-3’ (IDT Technologies, Iowa, United States) was used as a counterstain and targets the 16S rRNA of organisms in the domain of *Bacteria* [40]. Cryo-sectioned tissue sections were dehydrated in a graded ethanol series (70% and 80%) for 3 minutes each. Tissue sections were then immersed in a 0.2% Sudan Black solution (prepared in 96% ethanol) for 20 minutes and washed thrice with a 0.02% Tween solution (prepared in 1X PBS). A total of 25 μl of 25% formamide hybridization buffer (20 mM Tris-HCl [pH 8.0], 5M NaCl, 0.1% sodium dodecyl sulfate, and 25% formamide) containing 100 pmol of the labelled probe (50 μg/ml stock) was added to the sections and incubated overnight at 48°C. Slides were then immersed in 50 ml of wash buffer (0.5M EDTA and 5M NaCl, 20mM Tris-HCl [pH 8.0]) for 30 minutes in a 46°C water bath. After washing, slides were plunged into ice cold water for 5 seconds and left to dry.

### Confocal Laser Scanning Microscopy (CLSM)

Hybridized samples were mounted using Citifluor™ (Citifluor Ltd, Enfield Cloisters, London) and imaged using an Elyra PS.1 LSM780 inverted laser scanning confocal microscope (Carl Zeiss, Göttingen, Germany) fitted with a 63×/1.4 Plan-Apochromat oil immersion objective using the Zeiss Zen Black 2012 SP2 software suite. Laser power and gain were kept constant between experiments.

### Scanning Electron Microscopy

Excised skin samples were fixed using 2.5% glutaraldehyde (prepared in 0.1M PBS pH 7.4) for 48 hours at 4°C and then washed three times with 0.1M PBS. Fixed samples were then dehydrated with a graded ethanol series (once with 30%, 50%, 70%, 80%, 90% and twice with 100% for 15 minutes at each step) together with gentle agitation. Samples were then subjected to amyl acetate immersion for 30 minutes. Samples were next critical point dried with the Bal-Tec CPD-030 Critical Point Dryer (Bal-Tec AG, Balzers, Liechtenstein) overnight and deposited onto SEM specimen stubs using NEM Tape (Nisshin Em. Co. Ltd, Tokyo, Japan). Samples were then sputter coated with gold using a Bal-Tec SCD 005 sputter coater (Bal-Tec AG, Balzers, Liechtenstein). Samples were viewed using a JSM-6360LV (JEOL, Tokyo, Japan) scanning electron microscope.

### Cytokine Luminex MAP analysis

Luminex MAP analysis was performed using the Bio-Plex Pro™ Mouse Cytokine 23- plex Assay (Bio-Rad, California, USA) as previously described [41].

### Flow cytometry

Skin was cut into pieces and incubated in RPMI containing 10% serum, 0.2mg/ml Collagenase IV (Roche, Basel, Switzerland) and 20000U/ml of DNAse I (Roche, Basel, Switzerland) for 1 hour at 37°C. Cells were then passed through a 19 G syringe and filtered through a 100 μm cell strainer (BD Biosciences, New Jersey, USA) to obtain a homogenous cell suspension which was stained with the following fluorochrome or biotin-conjugated monoclonal antibodies (mAbs): mouse IA/IE (M5/114.15.2) (BD Biosciences, New Jersey, USA); Ly6G (1A8), CD64 (X54-5/7.1), F4/80 (BM8), EpCAM (G8.8) (Biolegend, San Diego, United States) and, CD45 (30F11), Ly6C (HK1.4), CD24 (M1/69), and CD11b (M1/70) (eBioscience, California, USA). Multi-parameter analyses of cell suspensions were performed on a LSR II (BD Biosciences, San Jose, USA). Data were analyzed with FlowJo software (TreeStar, Oregon, USA).

### Statistical Analyses

Statistical analyses were performed with GraphPad Prism software (Version 6.05 for Windows, California, United States). Comparison of weight and CFU titres were performed using the non-parametric Kruskal-Wallis test with Dunn’s multiple comparison post-test with p values <0.05 deemed significant. Cytokine and flow cytometry comparisons were performed using the Mann-Whitney U test. Principal component analysis was done in R (Version 3.3.2) with the packages factoextra (Version 1.0.4) and FactoMineR (Version 1.34).

### Ethics statement

All procedures were approved and performed in accordance with the Institutional Animal Care and Use Committee (IACUC) in Nanyang Technological University, School of Biological Sciences (ARF SBS/NIEA0198Z).

## RESULTS

### The minimum colonization dose for *E. faecalis* in wounds is 10 CFU

To investigate the colonization and infection dynamics of *E. faecalis* OG1RF in wounds, we first determined the colonization dose required to colonize 50% of excisional wounds in C57BL/6 mice (CD_50_). We examined a range of infection inocula, ranging from 10^1^ to 10^6^ CFU, at 24 hours post inoculation (hpi) and determined the CD_50_ to be 10^1^ CFU which resulted in 80% of the mice having recoverable CFU (Fig 1). The CD_90_ was 10^2^ CFU. In general, we observed that the median recoverable CFU for all inocula at 24 hpi was similar to the initial inoculum. At inocula of 10^1^ and 10^2^, we observed no visible, macroscopic signs of inflammation (**Supplementary Fig 1**). By contrast, at 24 hpi, inocula of 10^6^ resulted in visible, macroscopic inflammation with redness and accompanied by presence of serous exudates at the wound site of all infected mice (**Supplementary Fig 1**). Henceforth, we defined 10^6^ as the infectious dose (ID_90_). These findings suggest that the initial bacterial inoculum can result in wounds of two different states: colonization or infection.

**Figure 1:**
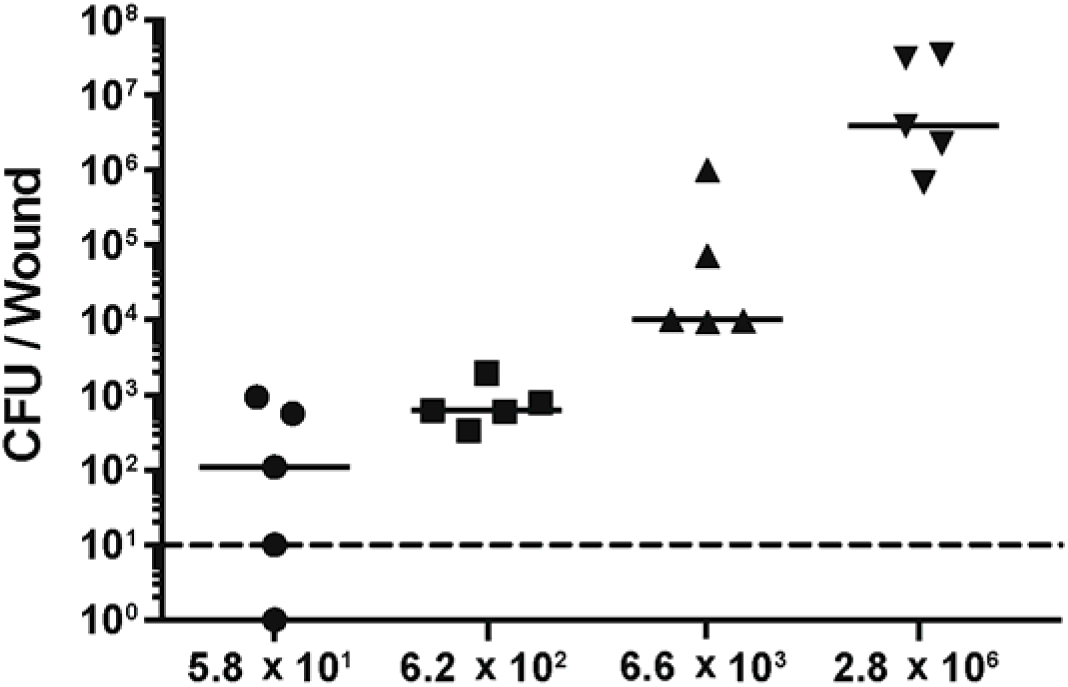
The CD_50_ of E. faecalis wound infection is 10^1^ CFU. Male C57BL/6 mice were wounded and infected with *E. faecalis* OG1RF with an inocula of 10^1^, 10^2^, 10^3^ or 10^6^ colony forming units (CFU). Wounds were harvested at 24 hpi and the recovered bacteria enumerated. Each dot represents one mouse, and the solid horizontal lines indicate the median. The horizontal dashed line indicates the limit of detection, n=5.

### *E. faecalis* infection is associated with high titer persistence in wounds

To further investigate the differences between low-inoculum colonization and high-inoculum infection dynamics, we inoculated mice with the CD_90_ colonization dose of 10^2^ CFU, or the infection dose of 10^6^ CFU, and monitored the mice for 7 days post inoculation (dpi). We observed that, regardless of the initial infection inoculum, viable bacteria were recovered at all time points (24 hpi to 7 dpi). However, after a 10^2^ CFU inoculation, *E. faecalis* persisted at 10^2^ CFU and only decreased at 7 dpi (Fig 2A). By contrast, when mice were inoculated with 10^6^ CFU, we observed a rapid increase to 10^8^ CFU by 8 hpi, followed by a decrease at 3 dpi to 10^5^ CFU which was maintained throughout the course of the experiment (Fig 2B). Consistent with this, at inocula of 10^6^, we observed visible inflammation and wound exudates only at 8 and 24 hpi, which resolved after 2 dpi (data not shown).

**Figure 2:**
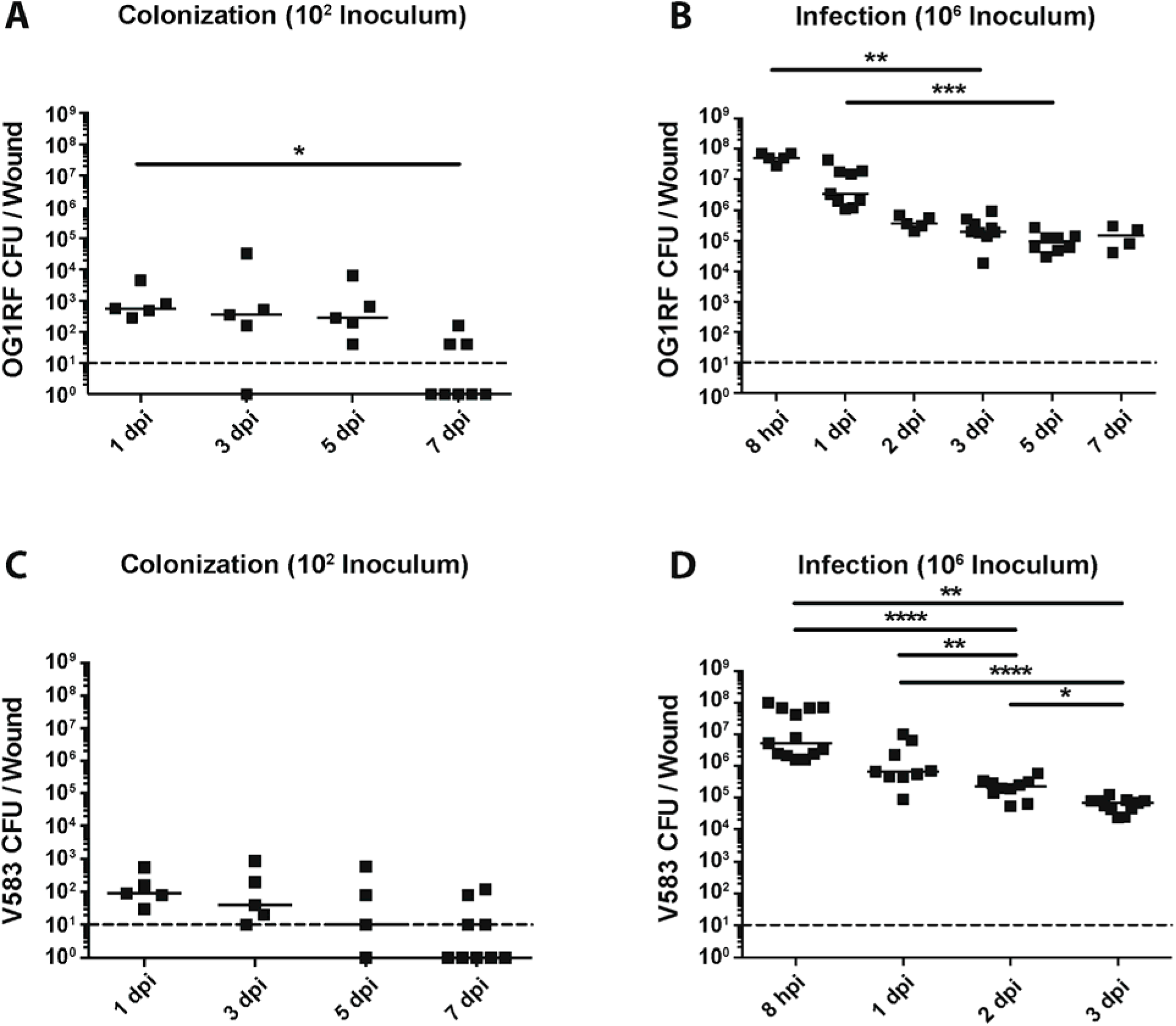
Colonization and infection dynamics of *E. faecalis* in wounds. Wounds were harvested at the indicated time points post-inoculation and the colony forming units (CFU) were enumerated. Mice were inoculated with **(A)** 10^2^ CFU of OG1RF, **(B)** 10^6^ CFU of OG1RF, **(C)** 10^2^ CFU of V583, or **(D)** 10^6^ CFU of V583. Each dot represents one mouse, and the solid horizontal lines indicate the median, N=2, n=≥5. Statistical analysis was performed using Kruskall-Wallis test with Dunn’s post-test to correct for multiple comparisons. * = p < 0.05, ** = p < 0.01.

*E. faecalis* wound infection dynamics were not strain-specific because the clinical blood isolate *E. faecalis* V583 [42] displayed similar infection kinetics to that of strain OG1RF (Fig 2 C,D). In addition, clinical *E. faecalis* wound isolates inoculated at the infection dose resulted in similar high titer wound infections at 8 hpi (**Supplementary Fig 2**). Together, these results demonstrate and confirm that the initial inoculum can determine the following states: colonization in the absence of increase *E. faecalis* titers and overt inflammation, or infection associated with acute bacterial replication and overt inflammation.

### MprF contributes to *E. faecalis* fitness during wound infection

To determine *E. faecalis* factors involved in wound colonization and infection, we examined the fitness of previously described biofilm factors as well as factors involved in immune defense. In competitive infections, we found that an Ebp null mutant was as fit as wild type OG1RF at 3 dpi, indicating that biofilm-associated Ebp was not important for wound infection (**Supplementary Fig 3**). Similarly, a Δ*srtAC* double mutant strain defective in biofilm formation, was not attenuated in fitness during coinfection (**Supplementary Fig 3**). Because we observed overt inflammation after high dose *E. faecalis* infection in wounds, we predicted that resistance to host immune killing may be important for its survival. *E. faecalis* encodes two paralogues of MprF [27, 28]. To address the contribution of these gene products to fitness in wounds, we co-infected wild type *E. faecalis* OG1X with the Δ*mprf1/2* strain, and found that while OG1X always outcompetes OG1RF to some degree (**Supplementary Fig 3**), the Δ*mprf1/2* mutant was massively outcompeted during co-infection (Fig 3). For wild type coinfection experiments, for reasons we do not yet understand, we observed bi-modality of OG1X out-competition of OG1RF, such that in each experiment some mice had much higher OG1X CFU than others. Together, these results suggest that traditional biofilm-associated factors may be less important for *E. faecalis* wound infection than its ability to resist immune defense mechanisms.

**Figure 3:**
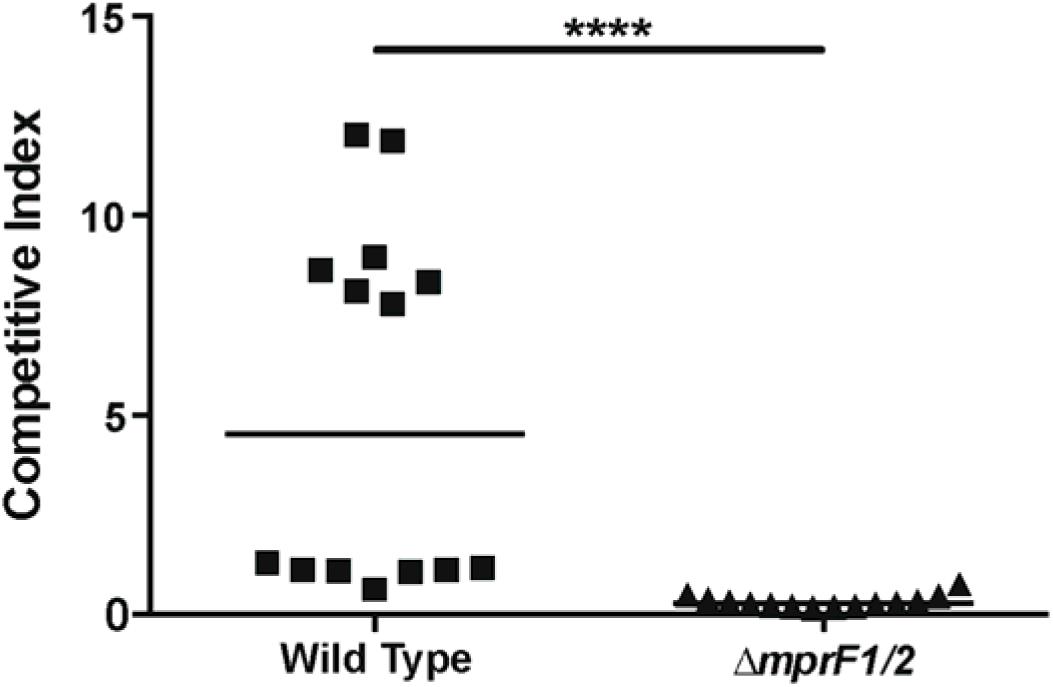
Multiple peptide resistance factor (MprF) contributes to fitness during *E. faecalis* wound infection. Male C57BL/6 mice were wounded and infected with a 1:1 ratio of *E. faecalis* strains OG1X:wild type OG1RF, or OG1X:OG1RF Δ*mprF1/2*, at 10^6^ CFU per inoculum. Wounds were harvested at 3 dpi and the recovered bacteria enumerated on selective media for each strain. Each dot represents one mouse, and the solid horizontal lines indicate the median, N=3, n=5. Statistical analysis was performed using Mann-Whitney U test. **** = p < 0.0001.

### *E. faecalis* forms microcolonies on the wound surface

Since we observed marked changes in the CFU recovered from infected wounds over time, we hypothesized that the spatial distribution of *E. faecalis* may also vary across time during infection. To address this question, we performed scanning electron microscopy (SEM) at 8 hpi and 3 dpi which represent the peak of infection and the onset of stable colonization, respectively. At 8 hpi, we observed *E. faecalis* microcolonies on infected wounds that appeared to be encased within a matrix, indicative of early biofilm development (Fig 4A). By contrast, at 3 dpi we were unable to detect *E. faecalis* on the surface of the wounds (Fig 4B). Since the CFU burden was still high at 3 dpi, we reasoned that *E. faecalis* may instead be embedded within the tissue. To determine the spatial localization of sub-superficial *E. faecalis* in 3 dpi wounds, we performed fluorescence *in-situ* hybridization (FISH) on 3 dpi wound samples. Using FISH probes specific for *E. faecalis*, we observed *E. faecalis* microcolonies at the wound edge (Fig 5A,C), and in the wound bed (Fig 5B,C). These results suggest that as *E. faecalis* wound infection proceeds, the bacteria first appear as biofilm-like microcolonies that are later encapsulated or internalized within the host tissues. We postulate that both properties may contribute to protection from the host immune response and persistence within wounds.

**Figure 4:**
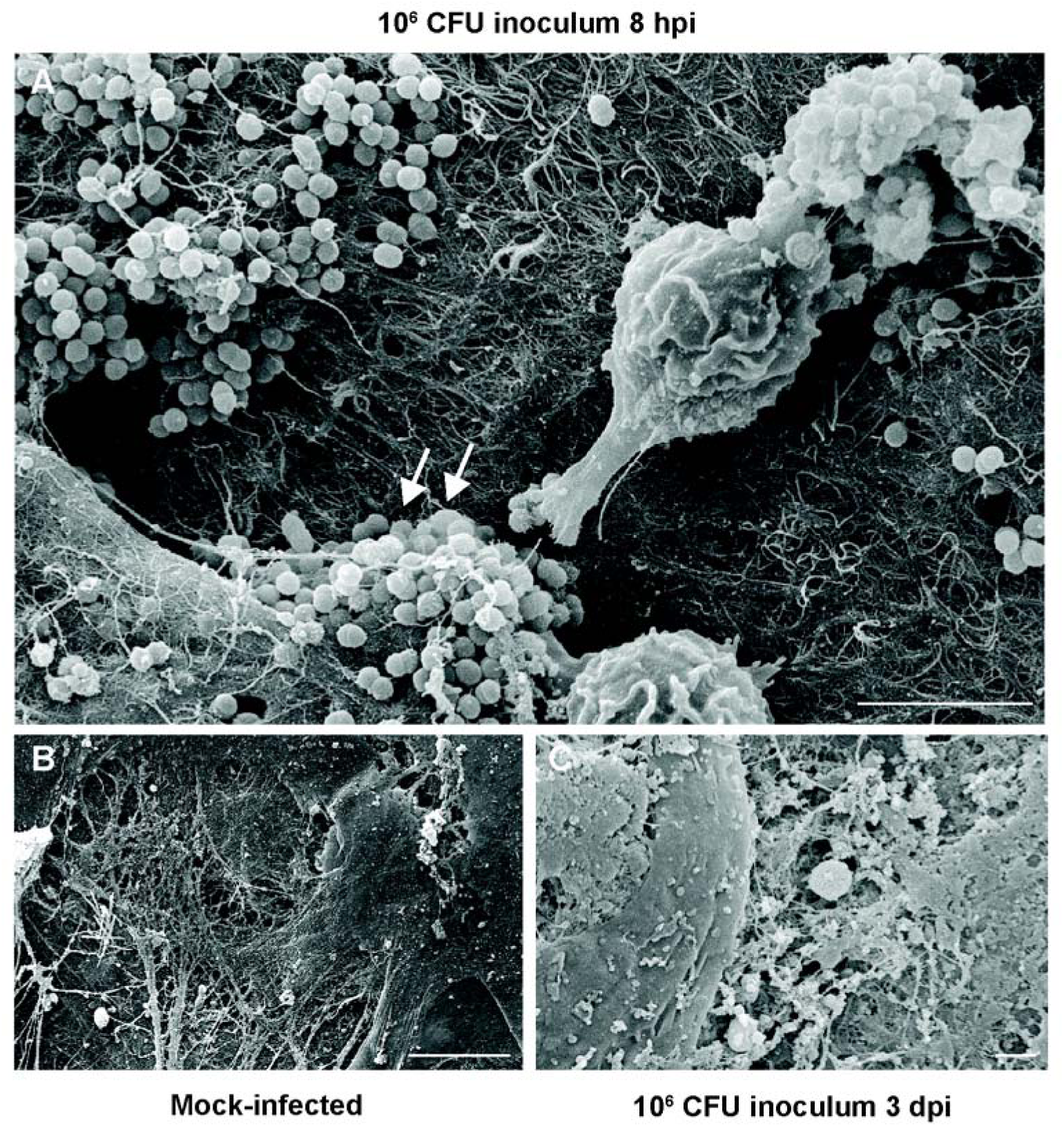
*E. faecalis* forms microcolonies in acutely infected wounds. Mice were wounded and infected with 10^6^ CFU *E. faecalis* OG1RF or mock-infected with PBS. Wounds were harvested at the indicated post-infection time points for scanning electron microscopy. *E. faecalis* microcolonies encapsulated by fibrous material were visible at 8 hpi (white arrows, **A**), but not in mock-infected wounds (**B**) or in infected wounds at 3 dpi (**C**). Bar represents 5 μm. Images shown are representative images from three independent experiments.

**Figure 5:**
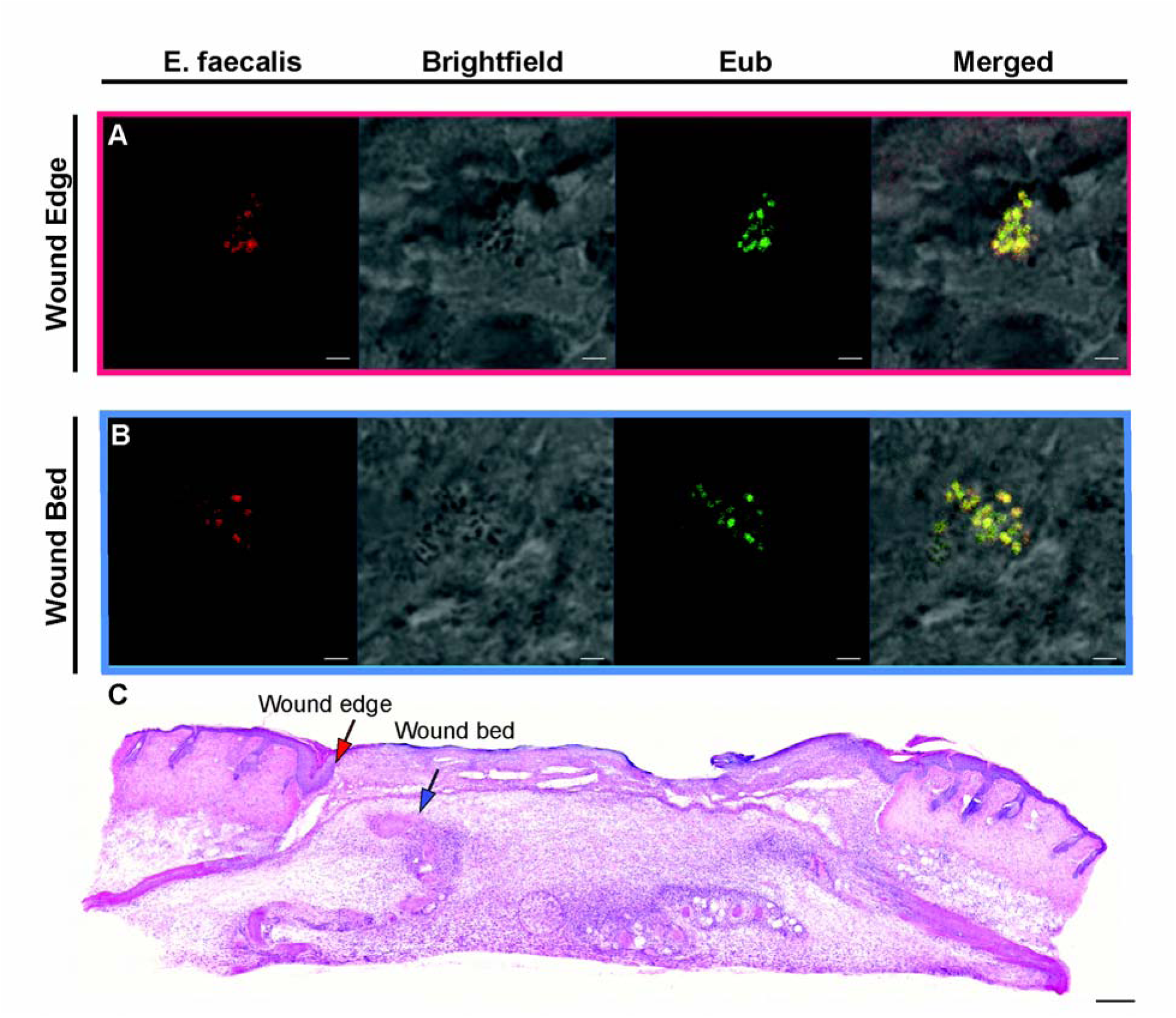
*E. faecalis* is present at the wound edge and in the wound bed. Male C57BL/6 mice were wounded and infected with 10^6^ CFU of *E. faecalis* OG1RF. Wounds were harvested at 3 days post-infection, cryosectioned, and subjected to (**A,B**) FISH or (**C**) H&E staining. (**A**) *E. faecalis* specific probes or probes specific for the domain bacteria (Eub) were used for FISH. The brightfield channel shows light microscopy images. Red and blue arrows (**C**) correspond to the red and blue boxes (**A,B**) and represent the wound edge and wound bed, respectively. (**A,B**) Bar represents 2 μm. Images shown are representative from three independent experiments.

### High titer *E. faecalis* infection alters wound healing and delays wound closure

Infection of wounds by *P. aeruginosa* and S. *aureus* correlate with delayed reepithelization and wound healing [43, 44]. To determine whether *E. faecalis* similarly affects wound healing, we performed histology on skin tissue obtained from wounds of infected mice at 7 dpi. Hemotoxylin and eosin (H&E) staining revealed a hyper-thickened epidermis, indicative of non-progressive wound healing, with delayed closure in the infected tissues which was not seen in the wounded, mock-infected controls (Fig 6A,B). Moreover, we also observed large numbers of polymorphonuclear leukocytes in H&E stained infected samples as late as 7 dpi as compared to mock-infected controls (Fig 6A). In addition, granulation tissue, which is indicative of dermal healing, was not properly formed in infected wounds, whereas healing was visible in mock-infected controls (Fig 6A,B). Long term persistence of *E. faecalis* also resulted in delayed wound closure (Fig 6C). These observations show that high titer *E. faecalis* infection negatively affects the wound healing process and delays the onset of wound closure.

**Figure 6:**
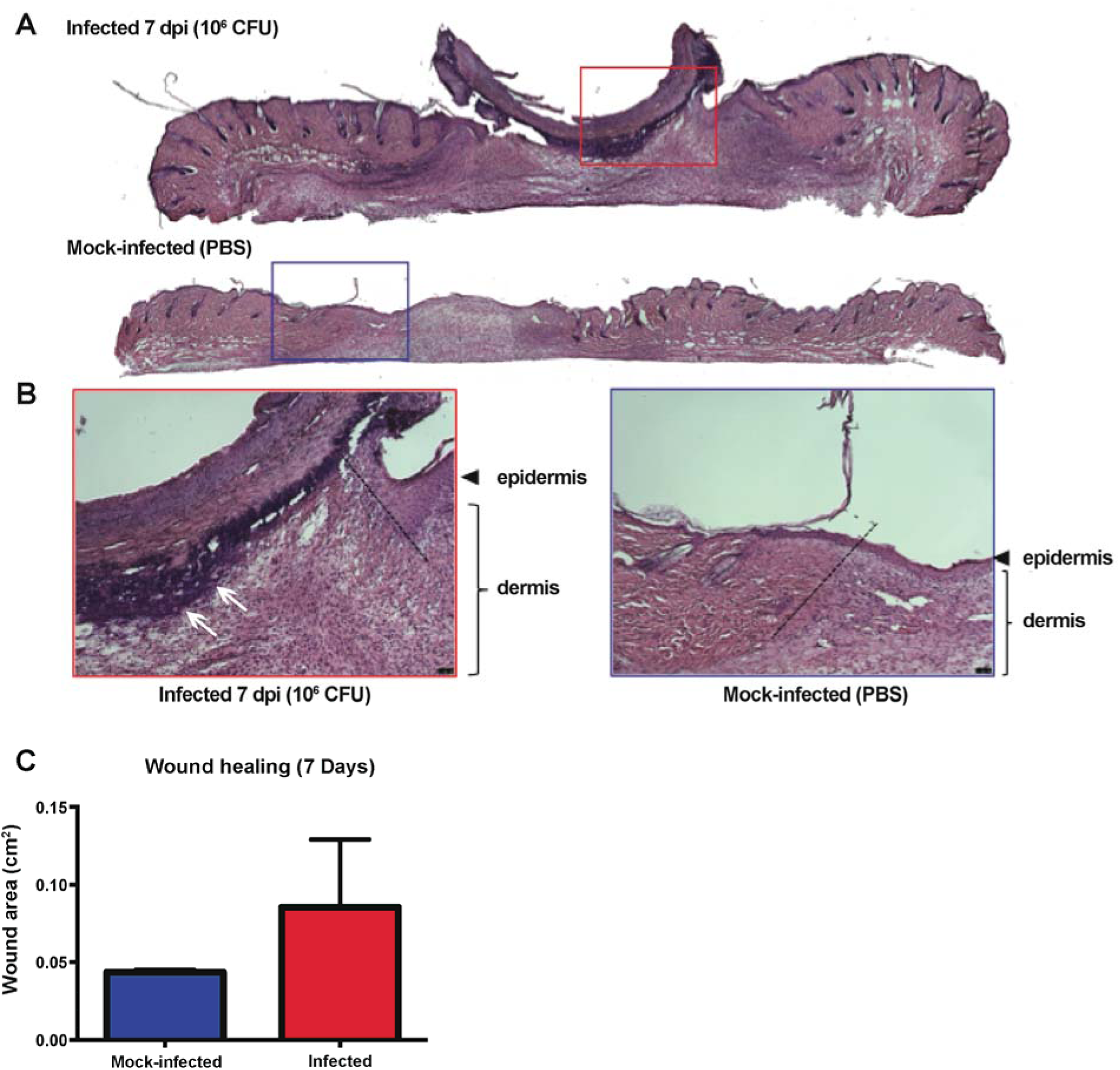
*E. faecalis* infection alters wound healing dynamics. Mice were wounded and infected as described above. Wounds were harvested at 7 dpi and subjected to H&E staining. (**A,B**) Red and blue boxes represent the wound edge for the infected wounds and mock-infected controls, respectively. (**B**) Higher magnification images of the boxes depicted in **A** and the dashed line indicate the wound edge. More clustered polymorphonuclear leukocytes are present at the 7 dpi wound (white arrows) but absent from the mock-infected wound. Bar represents 20 μm. Images are representative observations from three independent samples examined.

### *E. faecalis* can persist within wounds while escaping immune detection

We next hypothesized that *E. faecalis* might escape host detection during wound infection, contributing to its ability to persist and delay wound healing. Therefore, to examine the host immune response to *E. faecalis* infection, we first performed cytokine, growth factor and chemokine analysis on supernatants from wound homogenates inoculated with either 10^2^ or 10^6^ CFU, or PBS. At both 8 hpi and 3 dpi, wounds inoculated with 10^2^ *E. faecalis* CFU had cytokine and growth factor levels similar to the PBS controls (Fig 7A,B). By contrast, wounds infected with 10^6^ CFU displayed significantly higher levels of the inflammatory cytokine IL-1b, as well as growth factors/chemokines CSF3, CXCL1, CCL2, CCL3 and CCL4 compared to the controls at 8 hpi (Fig 7A**, Supplementary Fig 4A**), when macroscopic inflammation was observed. At 3 dpi, when *E. faecalis* wound titers resolved to 10^5^ CFU (Fig 2B), we observed significantly lower levels of IL-2, IL-5, IL-10, IL12-p70, CCL11, IFN-γ and CSF2 compared to both 10^2^ CFU-inoculated and PBS mock-infected wounds (Fig 7B**, Supplementary Fig 4B**). Reduced cytokine and chemokine levels during steady state infection suggest that *E. faecalis* may modulate the host immune response in wounds to promote persistence.

**Figure 7:**
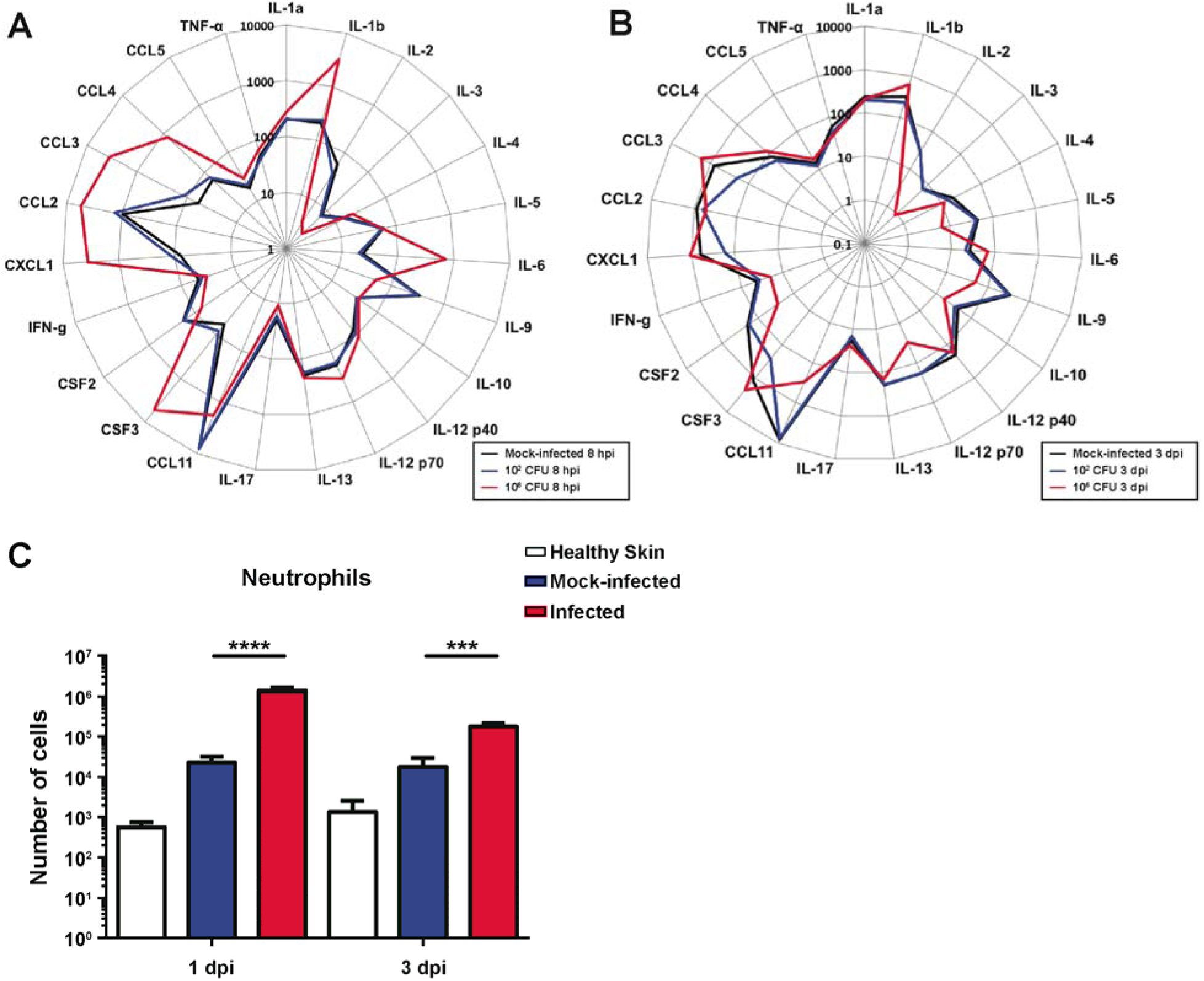
*E. faecalis* modulates the soluble and cellular host response at the wound site. Mice were wounded and infected with 10^2^ CFU or 10^6^ CFU of *E. faecalis*, or mock-infected with PBS. At the indicated times, wounds were processed into single cell suspensions and subjected to **(A-B)** cytokine analysis shown in pg/ml. **(C)** Total number of neutrophils (CD45+ MHCII^−^ CD11b^+^ Ly6G^+^) infiltrating into and accumulating in the skin analysed by flow cytometry during the immune response. N=2, n=5. Statistical analysis was performed using Mann-Whitney U test comparing infected against mock-infected wounds. * = p < 0.05, ** = p < 0.01, *** = p < 0.001, **** = p < 0.0001.

To gain further insight into the spectrum of soluble factors that were most associated with *E. faecalis* immune modulation during infection, we performed principal component analysis (PCA) (**Supplementary Fig 4C**). The PCA profiles of wounds infected with 10^6^ CFU at 8 hpi and 3 dpi were distinct and clustered separately, confirming that high inoculum infection results in a different inflammatory profile temporally (**Supplementary Fig 4C**). Differences in IL-1β, IL-2, IL-12p70 and CCL11 specifically explained the variation between the PCA profiles and best represented differences between all sample groups. Among these, IL-2, IL-12p70 and CCL11 were significantly decreased in the 10^6^ CFU infected group when compared to the mock-infected controls, suggesting that downregulation of these cytokines in particular may be associated with an attenuated immune response (**Supplementary Fig 4B**).

To complement the analysis of soluble immune effectors, we performed flow cytometry to quantify the immune cell types present at the wound sites. At 1 dpi, only neutrophil infiltration was significantly increased in the infected wound compared to mock-infected controls (Fig 7C). Increased neutrophil infiltration correlates with the upregulation of chemotatic chemokines at 8 hpi (Fig 7A). However, at 3 dpi, there were significantly more neutrophils, monocytes, macrophages, and monocyte derived cells in the infected wounds compared to and mock-infected wounds (**Supplementary Fig 6B**). Despite the presence of significant immune infiltrates at 3 dpi, the *E. faecalis* bacterial burden in the wounds persisted at >10^5^ CFU.

Taken together, these data demonstrate that high titer inocula, resulting in high titer wound infection, is associated with an acute inflammatory response concomitant with the peak of infection. The resolution of acute high titer infection to a lower steady state infection at 3 dpi corresponds both a suppression of cytokine and chemokine levels yet the presence of immune cellular infiltrate, suggests a complex immunomodulatory program that is insufficient to resolve acute *E. faecalis* wound infection.

## DISCUSSION

Globally, SSIs affect 7% and 15% of hospitalized individuals in developed and in developing countries, respectively [1]. SSIs can extend the average hospitalization of patients by 5 to 17 days [1]. Despite the prevalence and clinical importance of *E. faecalis* wound infections, we know nothing of its pathogenic mechanisms in this infection setting. Here, we established a modified mouse wound excisional model to study the infection dynamics of *E. faecalis* as a model for surgical site infections.

We show that acute high titer *E. faecalis* wound infection associated with ≥10^6^ CFU is associated with a robust cellular host immune response and visible signs of inflammation, along with delayed wound healing, whereas inflammation is suppressed or absent in lower titer infections. Our observations are consistent with reports showing that bacterial counts of ≥10^6^ perturbs healing in humans [45, 46]. However, despite an early robust inflammatory response, *E. faecalis* can persist in the local wound site regardless of the inoculum load.

Consistent with reports that most wound infections involve biofilms, we observe the presence of microcolonies at the surface of *E. faecalis* infected wounds at 8 hpi. However, a sortase null mutant, deficient in the surface display of a variety of biofilm-associated factors, was not attenuated in wounds. Together, these findings suggest that *E. faecalis* wound-associated microcolonies or biofilms require other bacterial or host factors for their development, and that currently understood biofilm factors are not required in this niche. Furthermore, we discovered that after 3 dpi, *E. faecalis* can be found at both the wound bed and at the epidermal wound edge, suggesting that *E. faecalis* reservoirs within host cells may promote persistence in this niche.

Importantly, we show that *E. faecalis* wound infection results in immunomodulation. At 3 dpi, IL-2, IL-12p70, and CCL11 levels were lower in infected wounds compared to mock-infected wounds, suggesting active immune suppression at the cytokine and chemokine level. However, we still observed significant immune cell infiltrate at 3 dpi, indicating that immune modulation may be insufficient to limit a full inflammatory response. Nevertheless, the pro-inflammatory cellular infiltrate was not able to clear *E. faecalis* from the wounds. Thus, it is tempting to speculate that *E. faecalis* wound infection includes an active immune evasion or immune suppression component, which contributes to high titer infection and long-term persistence, leading to the development of a chronic, non-healing wound. Further, even modest *E. faecalis-m*ediated immune suppression may provide an advantage for co-infecting organisms commonly found with *E. faecalis* in polymicrobial wound infections [10, 39]. Given the widespread prevalence of Enterococcal wound infections, further studies into factors that promote *E. faecalis* pathogenesis in wounds and its consequences on wound healing are critical.

## CORRESPONDENCE

Kimberly Kline, Singapore Centre for Environmental Life Sciences Engineering, Nanyang Technological University, SBS-B1n-27, 60 Nanyang Drive, Singapore 637551. telephone: +65 6592 7943, fax: +65 6316 7349

## ACKNOWLEDGEMENTS

We would like to thank Pei Yi Choo for her assistance with the FISH imaging. We would also like to thank Milton Kwek and Declan Lunny for their help and advice regarding the histology, sectioning, and staining of skin samples.

